# Vitamin D attenuates diabetic myocardial injury via the Erbb4/ferroptosis axis

**DOI:** 10.1101/2023.12.11.571190

**Authors:** Hanlu Song, Yufan Miao, Yujing Zhang, Luoya Zhang, Hao Chen, Lulu Tang, Wenjie Li, Chenxi Gu, Xing Li

## Abstract

**Background:** Hyperglycemia and hyperlipidemia lead to the ferroptosis, well as the phosphorylation of Erbb4, and thereby increase the risk of cardiac hypertrophy. Thus, our investigation aims to explore whether vitamin D could mitigate diabetic cardiac injury through modulation of the Erbb4/ferroptosis axis.

**Methods and results:** KKAy mice fed on a high-fat diet were utilized to construct the prediabetic model, which showed an up-regulated phosphorylation of Erbb4, with concurrent ferroptosis in cardiac tissues. Following the intervention with vitamin D for 16 weeks, the activity of Erbb4/YAP signaling was suppressed and the severeness of ferroptosis was improved. Meanwhile disturbances in glucose-lipid metabolism and insulin secretion induced by high fat were alleviated, along with improvements in cardiac hypertrophy and myocardial function. Moreover, we established an *in vitro* damage model by introducing H9c2 myocardial cells to high glucose (HG, 33.3 mM) and palmitic acid (PA, 0.25 mM). Unsurprisingly, similar results have been acquired after vitamin D supplementation. Subsequently, selective inhibitors of Erbb4 (Dacomitinib) and ferroptosis (Ferrostatin-1) were applied to evaluate the efficiency of Erbb4 signaling on modulating ferroptosis *in vitro*, and conclusively confirming that inhibiting of Erbb4 indeed reduce ferroptosis under HG and PA stimulus. Additionally, treatment of vitamin D was found to reduce cardiomyocyte hypertrophy and prevent cell death by inhibiting Erbb4 activity. Interestingly, the combined intervention of Vitamin D and Dacomitinib exerted a synergistic effect on ameliorating the abnormal conditions.

**Conclusions:** Our study unveils, the correlation between Erbb4 and ferroptosis in diabetic heart. Providing evidences that vitamin D supplementation can improve ferroptosis related diabetic cardiac injury through inactivation of Erbb4. Proposing that the combination treatment of vitamin D and Erbb4 inhibitors may emerge as a highly feasible clinical strategy for diabetic myocardial injury.

## Introduction

Hyperglycemia, insulin resistance and compensated hyperinsulinemia may predispose to cardiac dysfunction^[1]^. Consequently, individuals with diabetes are more susceptible to developing heart failure^[2]^. As a matter of fact, cardiac injury stands as the primary cause of death among diabetic patients, commonly presenting as myocardial hypertrophy, fibrosis that leading to diastolic dysfunction, and eventually ends up with systolic dysfunction^[3]^. However, there is no effective treatment available for diabetic cardiac injury so far.

*In vivo* studies have linked the presence of left ventricular hypertrophy and diastolic dysfunction to prediabetes^[4, 5]^. In contrast to the irreversible nature of diabetes, the abnormalities of prediabetes are highly likely to be reversible with appropriate and effective interventions, such as dietary behaviors. Therefore, it is reasonable to propose that intervening during the prediabetic phase might offer a more effective strategy to prevent the progression of cardiac damage toward heart failure. Vitamin D (VD) is a lipo-soluble vitamin known for its role in regulating blood glucose, improving insulin resistance, and more^[6]^. In the meantime. VD deficiency is intricately associated with the development of diabetic cardiac injury. Specifically, the vitamin D receptor (VDR), through its binding with active vitamin D, inhibits the nuclear translocation of FoxO1, thereby reducing autophagy and alleviating heart damage in individuals with diabetes^[7, 8]^. Furthermore, VD supplementation not only improves blood glucose and insulin levels but also alleviates diabetic cardiomyopathy by downregulating the AGE and hexosamine pathways, and reducing NF-kB activity in the cardiac tissue^[9]^. Nevertheless, there remains a scarcity of research that investigates whether vitamin D interventions during the prediabetic phase could effectively reverse myocardial injury.

Oxidative stress, as well as impaired antioxidant system under high glucose conditions form the pathogenic mechanism underlying diabetic cardiomyopathy^[10]^. In fact, a quantity of reactive oxygen species (ROS) and lipid peroxides are being secreted under the hyperglycaemic and hyperlipidaemic internal environment^[10, 11]^, inducing high levels of ferroptosis^[12]^, such the case of myocardial cells. The epidermal growth factor receptor (EGFR) family consists of receptor tyrosine kinases Erbb1-4, among which Erbb4 plays a crucial role in the process of cardiac hypertrophy and ventricular remodeling^[13, 14]^. Neuregulin-1 (NRG1), a member of the epidermal growth factor ligand family, is produced by vascular endothelial cells and capable of activating Erbb4 in cardiomyocytes with a paracrine manner^[13]^. Previous studies have indicated that elevated glucose levels increase NRG1 expression^[15]^, whereas intervention with vitamin D inhibited NRG1 expression while improving the hyperglycemic conditions. Proposing the idea that VD may suppress the activation of Erbb4 by reducing NRG1. Furthermore, Poursaitidis et al. found that cells with mutated EGFR were sensitive to ferroptosis^[16]^, while Wang et al showed that upregulation of Erbb2 protected cervical cancer cells from autophagy-induced ferroptosis^[17]^. Nevertheless, the definitive role of Erbb4 in regulating the incidence of ferroptosis in the heart remains uncertain.

Therefore, prediabetic model was produced by feeding KKAy mice with high-fat diet, and *in vitro* injury model was conducted on H9c2 cardiomyocytes with high glucose and palmitic acid. The current study was aimed to explore the impact of vitamin D on diabetic cardiac injury and its potential mechanisms, aiming to provide scientific evidence for targeted prevention and treatment strategies for diabetic cardiomyopathy.

## Materials and methods

### Chemical reagents and antibodies

VD_3_ was procured from Sigma Aldrich (St Louis, MO, USA). The cell culture medium (DMEM) was obtained from GIBCO (Thermo Fisher Scientific, Waltham, MA), and fetal bovine serum (FBS) was acquired from LONSERA (Lonsa scince srl, New Zealand, Australia). Ferrostatin-1(Fer-1) was purchased from GLPBIO (Glpbio, Montclair, USA), and Dacomitinib was purchased from MCE (MedChemExpress, NJ, USA). The sources of antibodies were as follows: NRG1(10527-1-AP), β-actin(20536-1-AP), SLC7A11(26864-1-AP) were purchased from Proteintech, Erbb4 (RT1276), YAP(RT1664), Phospho-YAP(ET1611-69), ACSL4 (ET7111-43), Ferritin (R1601-9), GPX4 (ET1706-45) were purchased from HUABIO. Phospho-Erbb4 (Tyr1284) (AF3445) were purchased from Affinity Biosciences.

### Animals and experimental design

All animal experiments were approved by the Life Sciences Ethics Review Committee of Zhengzhou University (Ethics No. ZZUIRB2021-GZR0141). Four-week-old specific pathogen free (SPF) male C57BL/6J mice and KKAy mice were purchased from Beijing Huafukang Biotechnology Co. The mice were housed in a barrier environment and acclimatized for 2 weeks, in which C57BL/6J mice were the normal control group (SC), and were fed basal chow (10 kcal% fat) and KKAy mice were fed high-fat chow (45 kcal% fat) for 6 weeks to construct a pre-diabetic model, and the criteria for successful modeling were: 6.7<FBG<16.7 mmol/L; 11.1<2h-BG<30 mmol/L. Successfully modeled KKAy mice were randomly divided into four groups according to blood glucose levels: model group (MSC), low-dose vitamin D intervention group (LVD, 0.42 IU/g/w), medium-dose vitamin D intervention group (MVD, 1.68 IU/g/w), and high-dose vitamin D intervention group (HVD, 4.20 IU/g/w). After 16 weeks of vitamin D intervention, mice were anesthetized with sodium pentobarbital, and heart tissues and blood were immediately removed and preserved for later use.

### Cell cultures and treatment

H9c2 cells were obtained from the First Affiliated Hospital of Zhengzhou University and cultured in DMEM medium supplemented with 10% FBS and 1% penicillin/streptomycin at 37°C in humidified 5% CO_2_ conditions. The cells were used in the experiments once they reached 70-80% confluence. High glucose and palmitic acid medium (HGPA, 33.3 mM d-glucose, 0.25 mM PA) was used to construct injury models by culturing cells for 24 hours. Addition of 10 nM calcitriol and/or 73.7 nM Dacomitinib or 1 μM Ferrostatin-1 pretreated the cells for 2h while co-culturing the cells with HGPA for 24h.

### Echocardiographic measurement

After 16 weeks of vitamin D intervention, echocardiography was performed by using Vevo2100 imaging system. Parasternal short-axis M-mode echocardiographic images were obtained at papillary muscles level. The following parameters were recorded to reflect cardiac function: The following parameters were measured and calculated: left ventricular ejection fraction (EF), fractional shortening (FS), left ventricular mass, left ventricular volume at systole and diastole (LVVol;s and LVVol;d), interventricular septal thickness at systole and diastole (IVS;s and IVS;d), left ventricular internal dimension at systole and diastole (LVID;s and LVID;d), left ventricular posterior wall thickness at systole and diastole (LVPW;s and LVPW;d).

### Hematoxylin & eosin (HE) , wheat germ agglutinin (WGA) and DAB & Prussian blue staining

The hearts fixed in 4% paraformaldehyde underwent sequential steps of dehydration, clarification, wax dipping, and embedding. Subsequently, the paraffin-embedded heart tissues were sliced into 3-5 μm sections. These sections underwent xylene, anhydrous ethanol, and 75% alcohol soaking, followed by dewaxing and staining with hematoxylin and eosin (HE). Observation of the sections was conducted using a light microscope, and images were captured for subsequent analysis. For wheat germ agglutinin (WGA) staining, the sections underwent dewaxing, antigen retrieval, and incubation with primary and secondary antibodies. Simultaneously, the WGA working solution was drop-stained, nuclei were restained, tissue autofluorescence was quenched, and the sections were examined using a fluorescence microscope, with images captured for analysis. The cross-sectional area of cardiomyocytes was measured using Image J software, and the resulting data were analyzed. Furthermore, sections were deparaffinized, stained with Prussian blue and DAB, and images were observed and captured using a light microscope.

### Oil red O and reactive oxygen species (ROS) staining

The tissues preserved at -80°C were embedded using OCT compound to prepare frozen sections. Subsequently, these sections were stained with oil red O following the provided guidelines, and the resulting images were observed and captured using a light microscope. For further analysis, the frozen sections were allowed to thaw to room temperature, ensuring moisture control and subsequent drying. Tissue autofluorescence was minimized, and a staining solution for reactive oxygen species (ROS) was meticulously added to the sections. After restaining the nuclei with DAPI, the sections were sealed and observed using fluorescence microscopy, with images captured for subsequent analysis.

### Serum biochemistry

FBG, total cholesterol (TC), triglyceride (TG), low-density and high-density lipoprotein cholesterol (LDL-C and HDL-C), creatine kinase (CK) and lactate dehydrogenase (LDH) levels were measured using the Nanjing JianCheng Assay Kits. Fasting serum insulin (ins) and 25(OH)D_3_ and were measured using Mlbio Enzyme-linked immunosorbent assay Kits.

### Measurement of Tissue Iron (Fe), glutathione (GSH) and malondialdehyde (MDA) levels

Heart tissue (50 mg) was precisely weighed and placed into a grinding tube containing grinding beads. Subsequently, 450 mL of physiological saline was added, and the tissue was thoroughly ground using a grinder. The resulting mixture was then centrifuged in a pre-cooled centrifuge for 10 minutes, after which the supernatant was collected for measurement. The protein concentration was determined, and Fe, GSH, and MDA levels were measured as per the provided instructions. Subsequently, values were calculated and analyzed using the designated formulas.

### Cell viability assay

Cells were seeded in 96-well plates, allowing them to adhere before the addition of medium containing various concentrations of intervening substances. Subsequently, 10 μL of Cell Counting Kit-8 (ApexBio, Houston, USA) solution was introduced into each well, and the plates were incubated at 37°C in an incubator for 1-4 hours. Absorbance at 450 nm was measured, and cell viability was calculated based on the recorded measurements.

### Detection of reactive oxygen species (ROS) in cells

Add a minimum of 1 mL of 10 μM DCFH-DA to the cultured cells, enabling its free passage through the cell membrane. Subsequently, intracellular lipase hydrolyzes DCFH-DA into DCFH, which is unable to traverse the cell membrane. Active oxygen within the cell oxidizes DCFH to form DCF, which exhibits fluorescence. The fluorescence intensity of DCF serves as an indicator to measure intracellular active oxygen levels. Incubate the cells for 20 minutes at 37°C in the cell incubator to facilitate the process. Remove any residual DCFH-DA that hasn’t entered the cells, then observe and capture images using a fluorescence microscope.

### Western blot analysis

Proteins extracted from murine hearts and cells were initially separated using SDS-PAGE gel and subsequently transferred onto polyvinylidene fluoride membranes. Following this, the membranes were blocked using either 5% skim milk or bovine serum albumin. Later, they underwent incubation with primary and secondary antibodies and were detected using an ECL detection system.

### Statistical analysis

The obtained results underwent statistical analysis via SPSS 25.0 software. For quantitative data, normality and tests for normality and variance homogeneity were performed. Quantitative data were presented as mean ± standard deviation. Between-group comparisons were performed using ANOVA for one-way multilevel quantitative data and ANOVA for repeated measures across multiple time points. Additionally, the LSD-t test was employed for two-way comparisons between groups, *Р*<0.05 was considered statistically significant.

## Result

### Vitamin D attenuates disorders of glucolipid metabolism and insulin secretion in diabetic mice

To explore the potential of vitamin D intervention in preventing diabetic myocardial injury, we induced prediabetes in KKAy mice and continued VD for intervention for 16 weeks. In comparison to the normal control group of C57BL/6J mice, these mice exhibited accelerated weight gain (fig.1a), elevated fasting glucose levels (fig.1b), compromised glucose tolerance (fig.1c-d), hyperinsulinemia (fig.1f), insulin resistance (fig.1g-i), and hyperlipidemia (fig.1j-k), albeit without significant changes in high-density lipoprotein levels (fig.1l). After vitamin D intervention, improvements were observed in all previously mentioned indicators, except for changes in body weight.

**Fig.1.**
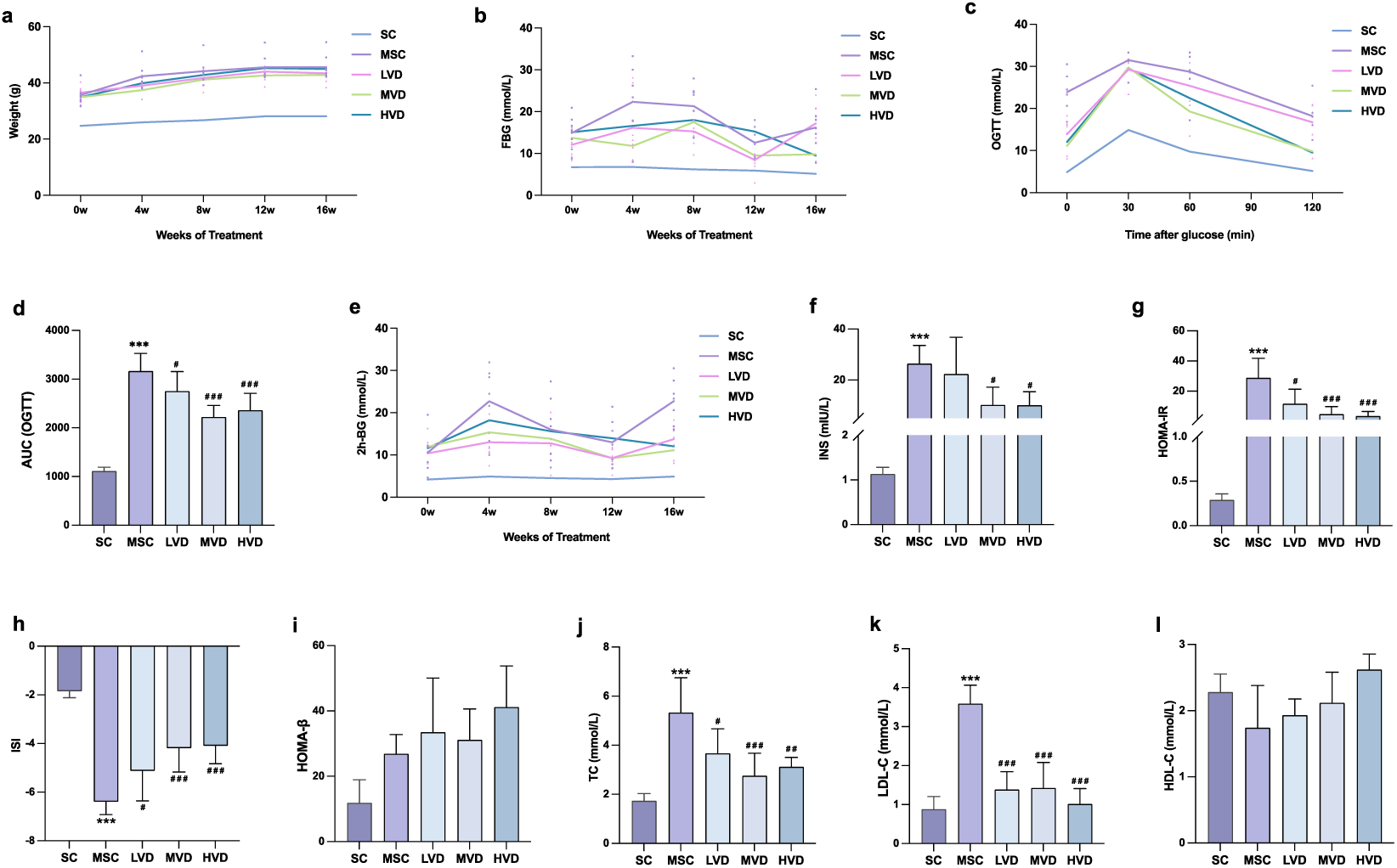
Vitamin D attenuates disorders of glucolipid metabolism and insulin secretion in diabetic mice. **a** Changes in body weight during vitamin D intervention. **b** Fasting blood glucose monitoring in five groups of mice in a 16-week VD intervention. **c-d** OGTT measurements and corresponding AUC following vitamin D intervention. **e** Monitoring of 2-hour postprandial blood glucose levels during vitamin D intervention. **f-i** Serum fasting insulin and respective insulin indices. **j-l** Total Triglyceride Levels and concentrations of high and low-density cholesterol measured using biochemical assays. Data were presented as Mean ± SD. \*\*\**P* < 0.001, compared to control group; **^#^***P* < 0.05, **^##^***P* < 0.01, **^###^***P* < 0.001, compared with model group.

### Vitamin D protects diabetic heart structure and function

Compared with SC mice, MSC mice had increased heart weight (fig.2a), cardiomyocyte hypertrophy, and markedly disorganised myocardial fibre structure with vacuoles (fig.2b). Additionally, the echocardiographic assessment revealed significantly increased LV mass (fig.2f-g), end-diastolic and end-systolic LV volumes (LV Vol;d and LV Vol;s, fig.2h-i), LV end-diastolic and end-systolic internal diameters (LVID;d and LVID;s, fig.2j-k) in MSC mice. Conversely, ejection fraction (EF%, fig.2d) and fractional shortening (FS%, fig.2e) were decreased. After the implementation of vitamin D intervention, the morphological structure of mouse cardiomyocytes showed tendencies toward intactness and relatively regular arrangement, with significantly improved cell hypertrophy and fiber breakage (fig.2b, n). Ventricular structure and function were also restored to varying degrees. Additionally, we assessed serum levels of 25(OH)D_3_, CK, and LDH (fig.2o-q). MSC mice exhibited lower 25(OH)D_3_ levels than SC mice, which increased after VD intervention. In contrast, levels of CK and LDH were higher in MSC mice compared to SC mice, but these levels decreased following VD intervention. Hence, vitamin D intervention can alleviate structural and functional changes in the hearts of diabetic mice.

**Fig.2.**
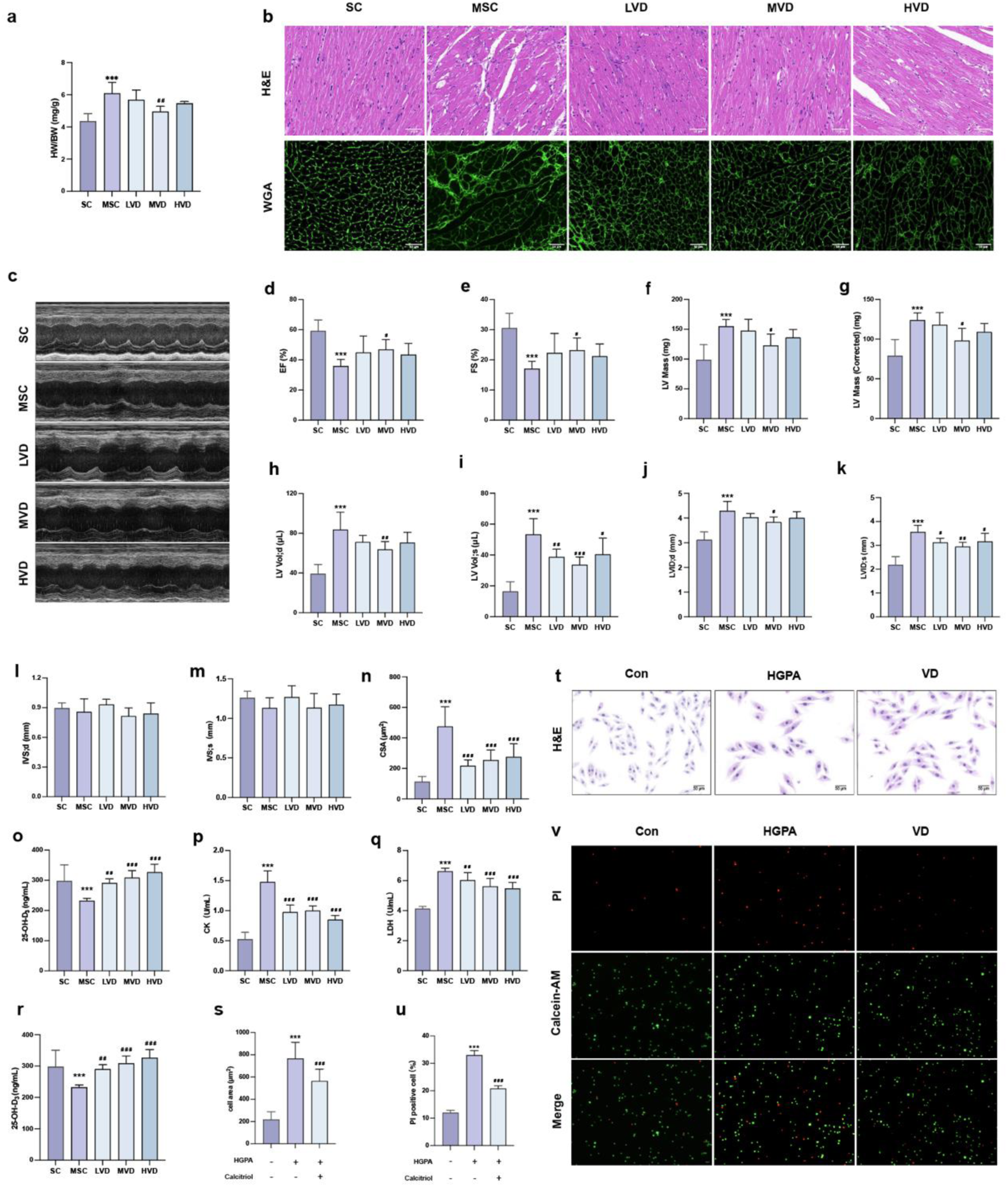
Vitamin D protects diabetic myocardial structure and function. **a** The heart weight to body weight was calculated. **b** Myocardial morphology was evaluated by H&E and wheat germ agglutinin (WGA) staining. **c-m** Echocardiographic parameters were evaluated: ejection fraction (EF%), fraction shortening (FS%), left ventricular (LV) mass, end-diastolic and end-systolic LV volumes (LV Vol;d and LV Vol;s), end-diastolic and end-systolic internal diameters (LVID;d and LVID;s), end-diastolic and end-systolic interventricular septal thickness (IVS;d and IVS;s). **n** Cross section area (CSA) of cardiomyocytes size was computed based on WGA staining. **o-q** Analysis of 25(OH)D_3_, CK and LDH levels in different groups of mice. **r** Cell viability analysis following various interventions. **s-v** Myocardial hypertrophy and cell death status were evaluated by images of H&E and PI/Calcein-AM staining. Data were presented as Mean ± SD. \*\*\**P* < 0.001, compared to control group; ^#^*P* < 0.05, ^##^*P* < 0.01, ^###^*P* < 0.001, compared with model group.

Furthermore, we initiated an in vitro model using H9c2 cardiomyocytes exposed to high glucose and palmitic acid to simulate cellular injury. The outcomes revealed that exposure to 33.3 mM HG + 0.25 mM PA for 24 hours (HGPA) led to reduced cell viability (fig.2r), increased cell death (fig.2u-v), and a noteworthy rise in the cross-sectional area of cardiomyocytes (fig.2s-t) when compared to the control group. We observed that pretreatment with calcitriol for 2 hours, followed by co-culturing with HGPA for 24 hours, significantly enhanced cell viability (fig.2r), reduced the number of deceased cells (fig.2u-v), and notably decreased the cross-sectional area (fig.2s-t) in the 10 nM calcitriol intervention (VD) compared to the HGPA group. In summary, these findings underscore the efficacy of vitamin D in ameliorating myocardial injury, both in vivo and in vitro.

### Vitamin D attenuates ferroptosis in heart tissue of diabetic mice

Diabetic mice manifest hyperglycemia and hyperlipidemia, conditions highly prone to triggering elevated levels of reactive oxygen species and lipid peroxides, resulting in the onset of ferroptosis. Our observations, conducted through DAB & Prussian blue staining alongside ROS staining, unveiled pronounced ferric hemoflavin-containing particles and a substantial accumulation of reactive oxygen species (ROS) within cardiac tissue slices from the model group mice (fig.3a-c). Notably, this condition exhibited improvement following VD intervention. Moreover, lipid deposition was notably reduced post-VD intervention, with only a few minute red lipid droplets observed in MVD heart tissue (fig.3a). In our investigation, we assessed the levels of markers associated with ferroptosis in the cardiac tissues of mice. The findings revealed significantly elevated levels of Fe and MDA (fig.3b,f) alongside notably reduced GSH levels (fig.3e) in MSC mice compared to SC mice. Additionally, cell death incidence was evaluated using the TUNEL assay (fig.3h), showing a reversal upon VD intervention. Western blot analysis (fig.3i-o) indicated a substantial increase in the relative expression of TFR1, ACSL4, NCOA4, and Ferritin in the cardiac tissues of MSC mice, which subsequently decreased following VD intervention. Conversely, the relative expression of SLC7A11 and GPX4 exhibited a significant decrease and increase, respectively, after VD intervention. These outcomes strongly suggest the occurrence of ferroptosis in the cardiac tissues of diabetic mice, a process mitigated by VD intervention.

**Fig.3.**
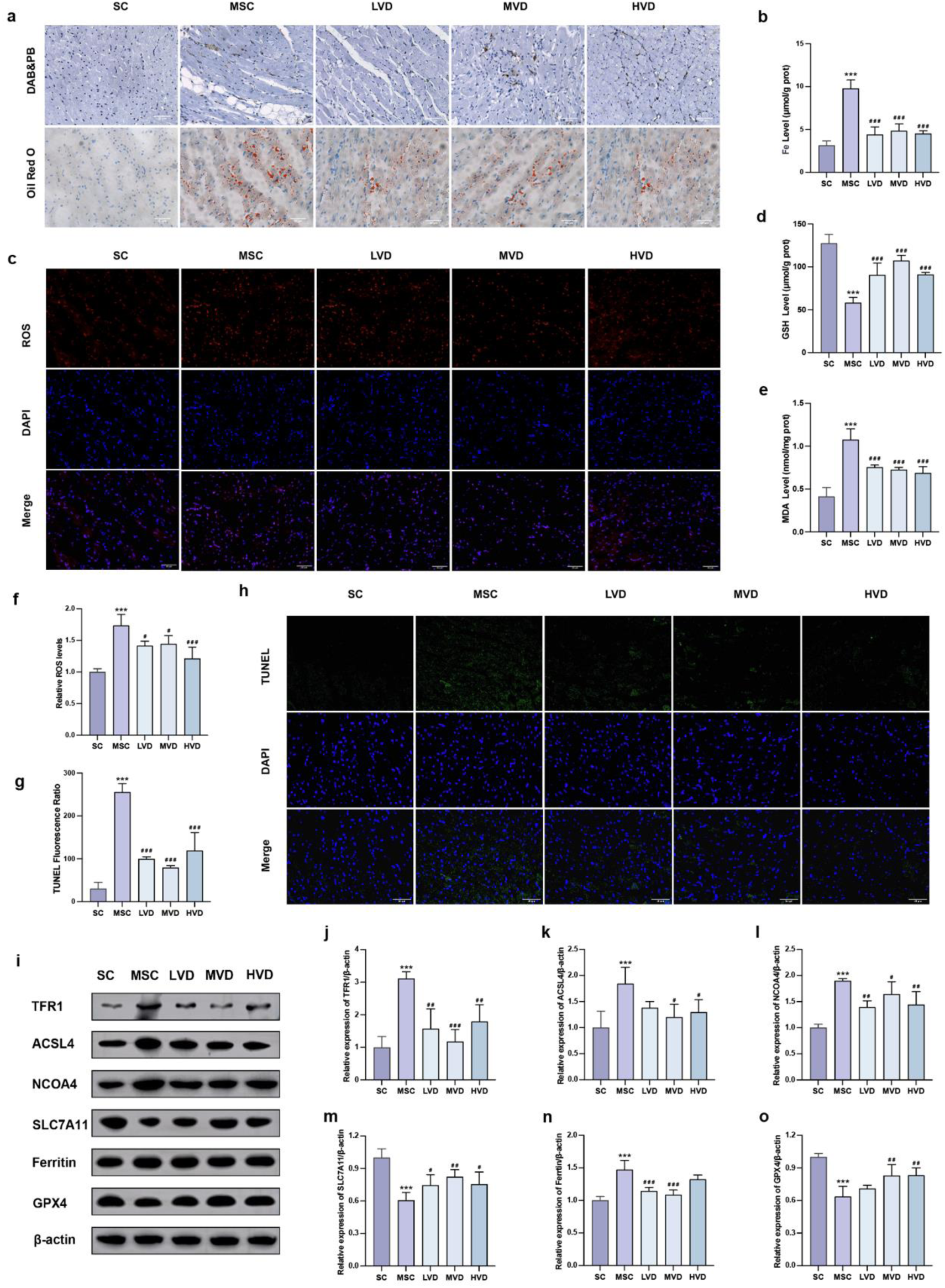
Vitamin D attenuates ferroptosis in heart tissue of diabetic mice. **a** Images of DAB & Prussian blue and Oil Red O staining in hearts from five groups of mice. **b-c** Representative fluorescence images and quantitative analysis of ROS measurement in mice. **d-f** Analysis of Fe^2+^, GSH and MDA levels in different groups of mice, n=6. **g-h** Quantitative analysis of TUNEL Fluorescence Ratio in myocardial tissues, Scale bar=50 µm, n = 3. **i-o** Representative Western blot images and quantitative analysis of TFR1, ACSL4, NCOA4, SLC7A11, Ferritin and GPX4 protein expressions in hearts of five groups, β-actin was used as a loading control. Data were presented as Mean ± SD, \*\*\**P* < 0.001, compared to control group; ^#^*P* < 0.05, ^##^*P* < 0.01, ^###^*P* < 0.001, compared with model group.

### Inhibition of ferroptosis attenuates high glucose and palmitic acid-induced cardiomyocyte injury

Subsequently, we further investigate whether inhibiting ferroptosis might alleviate cardiomyocyte injury induced by high glucose and palmitic acid (HGPA). *In vitro*, we initially exposed the cells to 1 μM Ferrostatin-1 for 2 hours, followed by co-incubation with HGPA for 24 hours (Fer-1) to inhibit ferroptosis. Subsequently, we observed a significant reduction in the cross-sectional area of Fer-1 cells compared to those solely exposed to HGPA (fig.4a-b). Additionally, the number of dead cells notably decreased (fig.4c-d), suggesting some degree of recovery from damage. Furthermore, we noticed elevated levels of Fe^2+^, MDA, and ROS, accompanied by a substantial decrease in GSH levels in HGPA cells compared to normal cells (fig.4e-i). Notably, these alterations were reversed upon application of the ferroptosis inhibitor. Subsequently, western blot analysis was performed to assess pivotal proteins implicated in ferroptosis. Results demonstrated a substantial upregulation in the relative expression levels of TFR1, ACSL4, and Ferritin, accompanied by a notable reduction in the relative expression of SLC7A11 and GPX4 subsequent to HGPA exposure (fig.4j-o). Intriguingly, these changes were reversed upon treatment with the ferroptosis inhibitor, Ferrostatin-1. This delineates the induction of ferroptosis in H9c2 cells by HGPA and highlights the potential of ferroptosis inhibition in mitigating HGPA-triggered cardiomyocyte injury.

**Fig.4.**
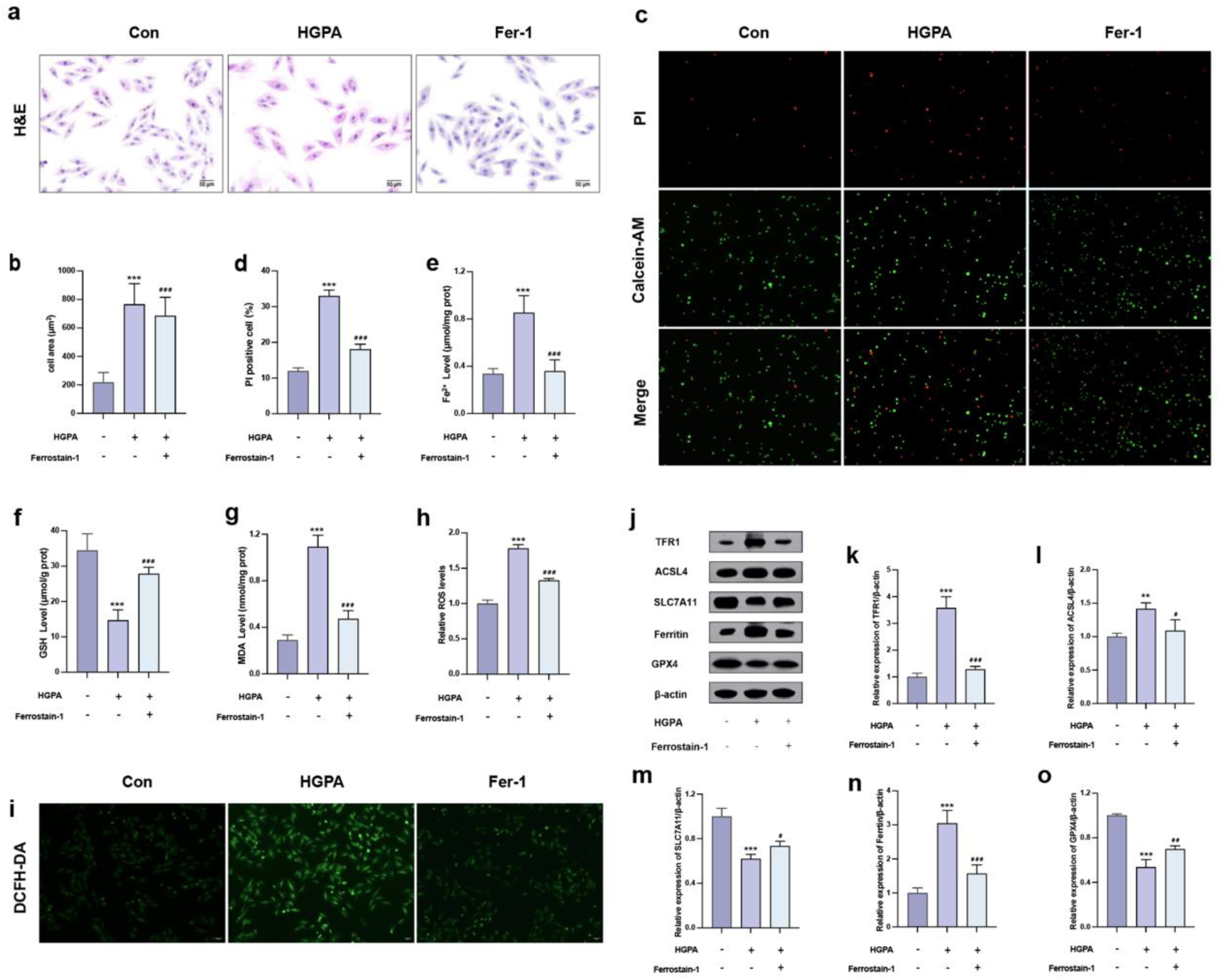
Inhibition of ferroptosis attenuates high glucose and palmitic acid-induced cardiomyocyte injury. **a-d** Myocardial hypertrophy and cell death status were assessed using images of H&E and PI/Calcein-AM staining. **e-g** Analysis of Fe, GSH and MDA levels in different groups of mice, n=6. **h-i** Representative fluorescence images and quantitative analysis of DCFH-DA staining for ROS measurement in H9c2 cells. **j-o** Protein expression of TFR1, ACSL4, SLC7A11, Ferritin and GPX4 in H9c2 cells. Data represent the results of three independent experiments and are shown as mean ± SD. \*\**P* < 0.01, \*\*\**P* < 0.001, compared to control group; ^#^*P* < 0.05, ^##^*P* < 0.01, ^###^*P* < 0.001, compared with model group.

### Vitamin D inhibits Erbb4 and YAP activation in cardiac tissues of diabetic mice

We observed significant cardiac hypertrophy in MSC mice (fig.2a-k), wherein Erbb4 and YAP have been established to play pivotal roles. Next, we explored the exact mechanism by which vitamin D. Therefore, we conducted a comparative analysis of the relative expression levels of pivotal proteins associated with potential pathways within the cardiac tissues of each murine cohort. The findings demonstrated a significant increase in NRG1 and phosphorylated Erbb4 expression levels within the cardiac tissues of MSC mice compared to SC mice (fig.5a-c). This upregulation was effectively mitigated following VD intervention. Additionally, there was a substantial reduction in phosphorylated YAP expression, correlating with inhibited YAP activation following VD intervention (fig.5d). This analysis indicates that vitamin D mitigates myocardial injury, potentially by downregulating the expression of NRG1 and inhibiting the activation of Erbb4 and YAP.

**Fig.5.**
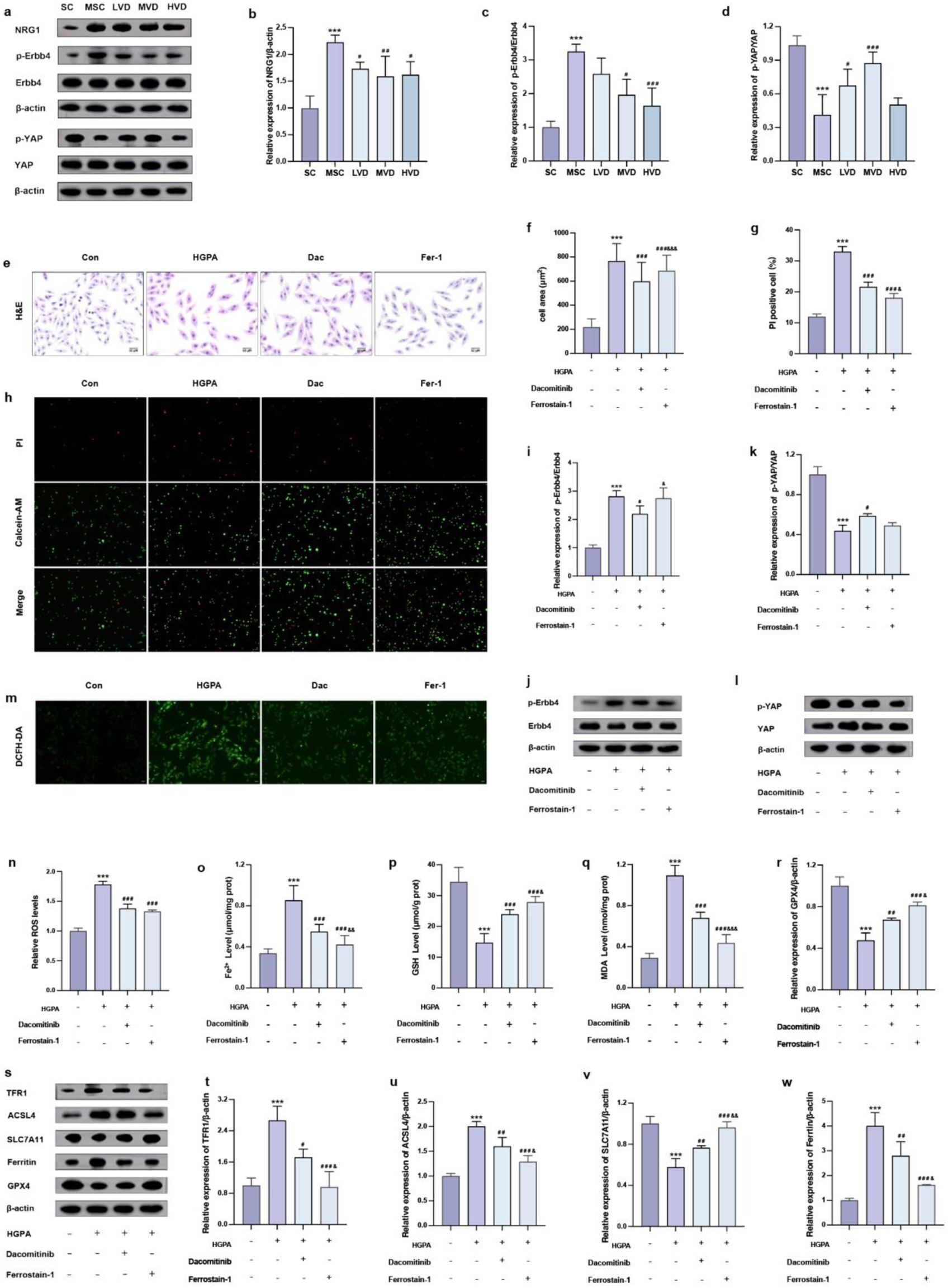
Vitamin D inhibits Erbb4 and YAP, Erbb4 inhibitors regulate YAP-mediated ferroptosis in H9c2 cardiomyocytes. **a-d** Representative Western blot images and quantitative analysis of NRG1, phosphorylated Erbb4, YAP and total Erbb4, YAP in the heart of mice, β-actin was used as a loading control. **e-h** Myocardial hypertrophy and cell death status were assessed by Images of H&E and PI staining. **i-l** Western blotting images of phosphorylated and total proteins of Erbb4 and YAP and values were presented by the ratio of phosphorylated expression to the total. **m-n** Representative fluorescence images and quantitative analysis of DCFH-DA staining for ROS measurement in H9c2 cells. **o-q** Analysis of Fe^2+^, GSH and MDA levels in H9c2 cells, n=6. **r-w** Protein expression of TFR1, ACSL4, SLC7A11, Ferritin and GPX4 in H9c2 cells. Data represent the results of three independent experiments and are shown as mean ± SD. \*\*\**P* < 0.001, compared to control group; ^#^*P* < 0.05, ^##^*P* < 0.01, ^###^*P* < 0.001, compared with model group; ^&^*P* < 0.05, ^&&^*P* < 0.01, ^&&&^*P* < 0.001, compared to Dac group.

### Erbb4 inhibitor modulates YAP-mediated ferritin deposition and hypertrophy in H9c2 cardiomyocytes

During the progression of diabetic myocardial injury, we observed the activation of Erbb4, simultaneous suppression of YAP activity, and the presence of ferroptosis. Next, we aim to explore the precise interrelation among these three factors. In our in vitro experiments, cells underwent pre-treatment 73.7 nM Dacomitinib for 2 hours, followed by a 24-hour co-culture with HGPA (Dac), aiming to inhibit Erbb4 phosphorylation and observe changes in other indicators. We found a decrease in the number of deceased cells (fig.5g-h) and a notable reduction in cell cross-sectional area (fig.5e-f) in Dac cells compared to HGPA cells. Additionally, we observed a decline in phosphorylated Erbb4 levels (fig.5i-j) coupled with a substantial increase in phosphorylated YAP levels (fig.5k-l), which corresponded to the inhibition of ferroptosis (fig.5m-w). Importantly, no significant difference was observed in p-Erbb4 and p-YAP levels between Fer-1 cells and Dac cells. Thus, our findings elucidate that Erbb4 functions upstream of YAP, and the diminished phosphorylation level thereof may attenuate YAP-mediated ferroptosis.

### Vitamin D inhibits Erbb4/YAP-mediated ferroptosis and rescues myocardial injury

Our previous in vivo study demonstrated a decrease in p-Erbb4 expression following vitamin D intervention. Subsequently, we explored whether vitamin D mitigates myocardial injury by modulating Erbb4/YAP-mediated ferroptosis. Therefore, in vitro experiments, we added 10 nM calcitriol and 73.7 nM Dacomitinib for 2h pretreatment followed by 24h co-culture with HGPA (VD+Dac). Compared with VD cells, VD+Dac cells showed a significant reduction in cell cross-sectional area (fig.6a-b) and a reduced number of dead cells (fig.6c-d). In addition, we observed a significant reduction in phosphorylated Erbb4 levels (fig.6e-f), a significant increase in phosphorylated YAP levels (fig.6g-h), and an inhibition of ferroptosis (fig.6i-s). The above analyses suggest that vitamin D inhibits Erbb4/YAP-mediated ferroptosis and rescues myocardial injury, and that Erbb4 inhibitors enhance the therapeutic effect of VD on myocardial injury.

**Fig.6.**
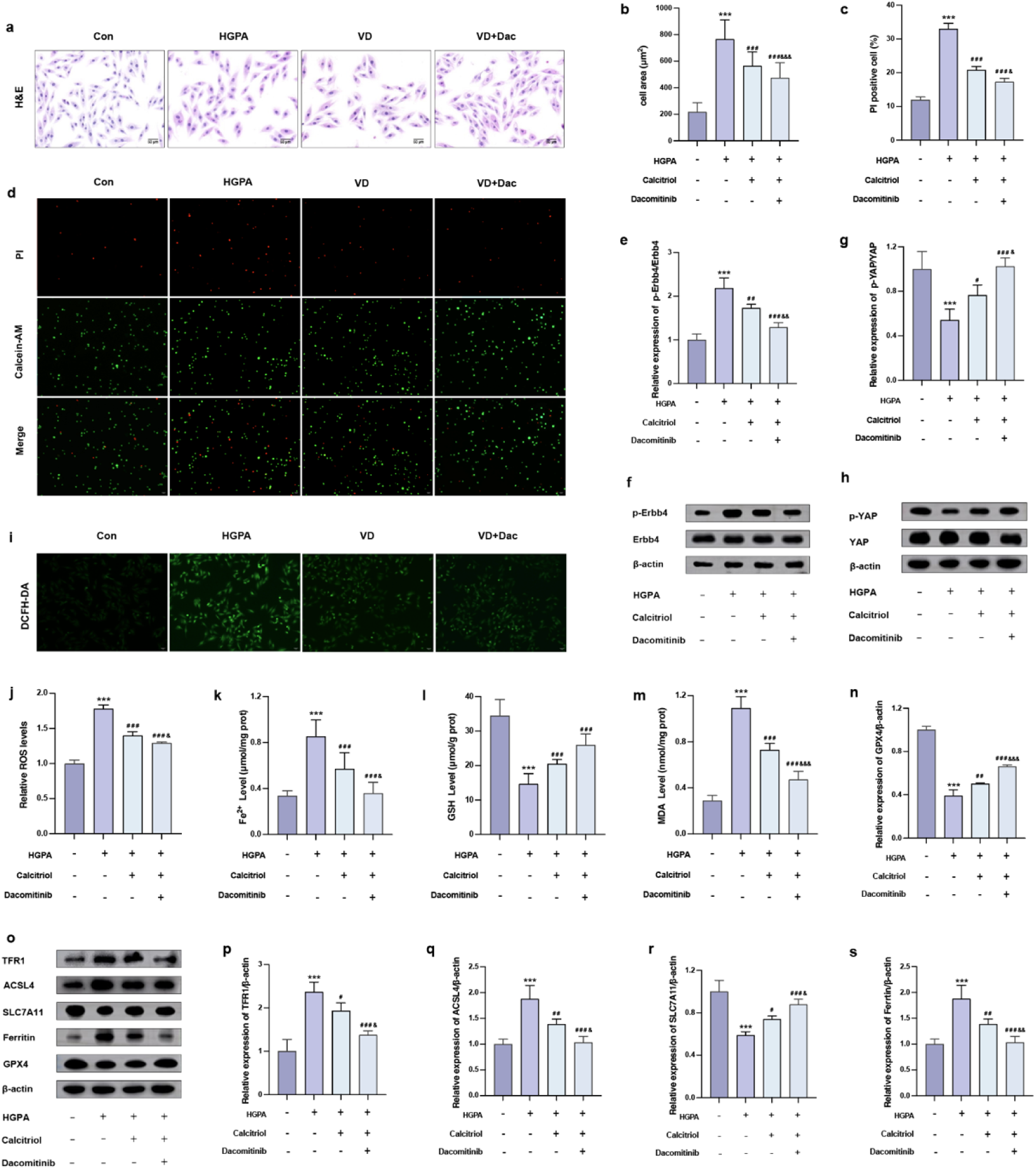
Vitamin D inhibits Erbb4/YAP-mediated ferroptosis and rescues myocardial injury. **a-d** Myocardial hypertrophy and cell death status were assessed by Images of H&E and PI staining. **e-h** Western blotting images of phosphorylated and total proteins of Erbb4 and YAP and values were presented by the ratio of phosphorylated expression to the total. **i-j** Representative fluorescence images and quantitative analysis of DCFH-DA staining for ROS measurement in H9c2 cells. **k-m** Analysis of Fe^2+^, GSH and MDA levels in H9c2 cells, n=6. **n-s** Protein expression of TFR1, ACSL4, SLC7A11, Ferritin and GPX4 in H9c2 cells. Data represent the results of three independent experiments and are shown as mean ± SD. \*\*\**P* < 0.001, compared to control group; ^#^*P* < 0.05, ^##^*P* < 0.01, ^###^*P* < 0.001, compared with model group; ^&^*P* < 0.05, ^&&^*P* < 0.01, ^&&&^*P* < 0.001, compared to VD group.

## Discussion

The primary findings of our study are as follows: 1) VD intervention during the pre-diabetic stage resulted in the attenuation of symptoms, such as disorders of glucose-lipid metabolism and insulin resistance, as well as varying degrees of ventricular paramorphia and dysfunction; 2) Diminished levels of Erbb4 phosphorylation effectively attenuated ferroptosis, and inhibited high glucose and palmitate-induced hypertrophy and death of H9c2 cardiomyocytes; 3) VD rescued myocardial injury through inhibition of the Erbb4/ferroptosis axis to rescue myocardial injury, whereas Erbb4 inhibitor showed a synergistic effect with VD on alleviating myocardial injury. To the best of our knowledge, this study reveals for the first time the relationship between Erbb4 and ferroptosis in diabetic cardiopathy, which provides novel evidence that VD can ameliorate diabetic cardiac injury, and that the combination of VD and Erbb4 inhibitor may be a feasible clinical strategy for treating diabetic myocardial injuries.

The effects of vitamin D on prediabetes present a contentious landscape in existing studies. Some studies suggest that VD supplementation during prediabetic stage with notably low VD levels improves insulin sensitivity, diminishes the risk of diabetes development, and improves insulin resistance in prediabetic rodents, while others have shown that treatment of VD_3_ for 24 consecutive months does not improve OGTT-derived indices of β-cell function in prediabetic patients^[18–20]^. Correspondingly, KKAy mice represent a spontaneously diabetic model characterized by hyperglycemia, impaired glucose tolerance, insulin resistance, and obesity induced by a high-fat diet. In our study, VD intervention for 16-week commencing from the prediabetic stage effectively mitigated disturbances in glucose-lipid metabolism and insulin resistance, along with improvements in cardiac function and ventricular structure of KKAy mice. In both prediabetic and diabetic patients, serum 25(OH)D levels exhibit a negative correlation with the risk of cardiovascular disease^[21]^, and VD is associated with the cardiac autonomic function and metabolic status in individuals with prediabetes^[22]^. However, the findings were all derived from observational studies and could not reveal a direct causal link between VD status and cardiac structure and function. Besides, the results from several interventional trials corroborate with our conclusions, indicating that VD supplementation has a modest beneficial impact on atherosclerotic cardiovascular disease risk in individuals with prediabetes^[23]^. Additionally, treatment with high-dose VD in VD-deficient prediabetic men led to a reduction in the increase of left atrial volume^[24]^.

In the meanwhile, MSC mice displayed symptoms of hyperglycemia and hyperlipidemia, owing to increased reactive oxygen species (ROS) production within the cytoplasm. This pathological process led to reduced cell viability and the accumulation of lipid peroxides^[25, 26]^, both known risk factors of ferroptosis. Recently, ferroptosis has emerged as a novel therapeutic target for cardiovascular diseases^[27]^. Furthermore, it has also been documented that ferroptosis plays a crucial role in the pathogenesis of diabetic cardiomyopathy^[28]^. Based on our results, inhibition of ferroptosis with Fer-1, a ferroptosis inhibitor, was shown to attenuate HGPA-induced death and hypertrophy of H9c2 cardiomyocytes; accordingly, key effector proteins associated with ferroptosis exhibited significant alterations in cardiac tissues, which was partially mitigated by VD intervention. Previously, VD/VDR has been demonstrated to avert ferroptosis in hippocampal neuron via activating the Nrf2/HO-1 signaling^[29]^. Additionally, VD has been shown to inhibit ferroptosis in liver, kidney, intestine, and hypoxic-ischemic brain tissues through various pathways^[30–33]^, which is consistent with our observations.

We found *in vivo* that vitamin D alleviates myocardial injury, likely by downregulating NRG1 expression and inhibiting Erbb4 activation. Erbb4, primarily situated on the cardiac cell membrane, belongs to the epidermal growth factor receptor (EGFR) family^[34]^, which possesses a growth-inhibitory property and is crucial for the typical development and sustenance of the heart, mammary gland, and nervous system^[34–36]^. Emerging researches have underscored the pivotal role of Erbb4 in the genesis of heart-related pathologies. For instance, tissue specific deficiency of Erbb4 in cardiac endothelial cells has been demonstrated to mitigate myocardial hypertrophy and fibrosis in the context of cardiac stress injury^[14]^. Yet selective activation of Erbb4 has been illustrated to induce cardiomyocyte hypertrophy^[13]^. Whereas mice with a mutational deletion on *Nrg1* or its corresponding receptor Erbb4 exhibited arrested development of ventricular myocardial trabeculae and succumbed during mid-embryogenesis^[37]^; and systemic blockade of Erbb4 function in mice modeling myocardial infarction intensified cardiac fibroblast senescence and apoptosis, consequently exacerbating inflammation^[38]^. These inconsistent results prompted us to presume that cardiac Erbb4 functions differently with upgrowth and aging.

Intriguingly, Li et al found that patients with diabetic cardiomyopathy suffered impaired NGR1/Erbb signaling, and aerobic exercise can elevate NRG1 expression and activate Erbb4, thus fostering cardiac repair by stimulating endogenous regeneration^[39]^, which is contrary to our findings. While other reports have proposed the possibility that in non-ischemic heart failure, the endothelial microvascular system exposed to oxidative stress exhibits a compensatory effect by increasing cardioprotective NRG1-β^[40]^ . Furthermore, studies by breast cancer researchers have shown that hyperglycemia induces an upregulation of NRG1^[15]^, which supports our observations that NGR1 experienced a compensatory increase during the pre-diabetic phase of cardiac injury, which subsequently induces the activation of Erbb4 on cardiomyocytes, and promoting myocardial hypertrophy. We found that vitamin D can inhibit the activation of Erbb4, which is in consistency with the results of several past studies, showing a significant role of vitamin D in glycemic control. We therefore have substantiated a possibility: vitamin D improves the blood glucose levels of diabetic mice, subsequently attenuating the upregulation of NRG1 induced by high blood sugar, thereby inhibiting the activation of Erbb4, and alleviating myocardial injury.

In our investigation, we observed the activation of yes-associated protein 1 (YAP), a nuclear effector within the Hippo pathway, in the hearts of the model mice. Previous research has elucidated the role of YAP in mediating compensatory cardiac hypertrophy during acute pressure overload^[41]^. Moreover, this mechanism is believed to contribute, to some extent, to the development of dilated cardiac hypertrophy and pathological hypertrophy^[42, 43]^. MSC mice exhibited a noteworthy increase in left ventricular volume and internal diameter, indicative of dilated cardiac hypertrophy, wherein YAP activation appeared to be involved. Simultaneously, we observed that the activation of YAP was inhibited following VD intervention. The results were in accordance with the findings of some other studies support, indicating that co-administration of VD with Aflatoxin B1 intercepts the Hippo pathway^[44]^. Additionally, Vitamin D and its analogs have been shown to promote wound healing and inhibit inflammatory responses via downregulation of TGF-β-mediated YAP phosphorylation^[45, 46]^.

MSC mice in our experiment showed upregulated levels of Erbb4 activation, concurrent suppression of (p)YAP activity, and the presence of more ferroptosis in the cardiac tissues. Previous studies have highlighted the susceptibility of EGFR mutant cells to ferroptosis^[46]^, while upregulation of Erbb2 has been demonstrated to generate protective effect on cervical cancer cells from autophagy-mediated ferroptosis^[16, 17]^. Considering these insights, we hypothesized that Erbb4 signaling might regulate ferroptosis during diabetic heart injury. To investigate their interrelations, in vitro experiments utilizing Erbb4 and ferroptosis inhibitors have been conducted on HGPA treated H9c2 cardiomyocytes. Once being phosphorylated, the intramembrane protein Erbb4 hydrolysis and releases the soluble intracellular domain (ICD) into the nuclear for gene transcription^[47]^. Specifically, the PPxY motifs facilitate its interaction with the WW structural domains in YAP, consequently enhancing its transcriptional activity^[47]^. Furthermore, numerous in vitro studies have highlighted YAP’s ability in modulating ferroptosis through diverse mechanisms^[48–50]^. As anticipated, our findings suggest that Erbb4 operates as an upstream regulator of YAP, whose phosphorylation could impede the processing of YAP-mediated ferroptosis. Furthermore, our animal experiments illustrated a decrease in p-Erbb4 expression following vitamin D intervention, aligning with the effects exerted by Erbb4 inhibitors. Notably, the combined intervention of vitamin D and Erbb4 inhibitor demonstrated heightened efficacy in alleviating myocardial injury. Additionally, vitamin D exhibited the capability to inhibit Erbb4/YAP-mediated ferroptosis, which eventually rescued myocardial injury.

In summary, vitamin D plays a beneficial role in mitigating diabetic heart injury. Nonetheless, our study has certain limitations. While supporting the feasibility of this theory, the complex in vivo environment entails numerous influencing factors. Hence, further validation in the population or in Erbb4 knockdown/out animal models is essential to consolidate this mechanism. Our study primarily emphasizes the role of vitamin D in reducing cardiac hypertrophy. Future investigations could delve deeper into exploring its pharmacological impact on inflammation, fibrosis, or diastolic dysfunction.

## ACKNOWLEDGMENTS

We would like to thank the members of the Li labs for helpful discussion.

## AUTHOR CONTRIBUTIONS

H.Song and Y.Miao performed the experiments. H.Song analyze data. H.Chen and L.Tang assisted the animal experiments. H.Song, Y.Zhang and Y.Miao designed project. H.Song and L.Zhang wrote the manuscript. X.Li, Y.Zhang, C.Gu and W.Li supervised experiments and critically reviewed the manuscript. All authors read and approved the final manuscript.

